# Overview of non-coding RNAs with CAG repeats and the case of mutation-containing circRNA in polyglutamine disease patients

**DOI:** 10.1101/2025.08.13.669846

**Authors:** Weronika Pawlik, Magdalena Woźna-Wysocka, Magdalena Jazurek-Ciesiołka, Jarosław Dulski, Jarosław Sławek, Edyta Kościańska, Tomasz Witkoś, Agnieszka Fiszer

## Abstract

CAG trinucleotide repeats are present in protein-coding and non-coding RNAs (ncRNAs). They are polymorphic in length and can cause neurodegenerative diseases when expanded, e.g., spinocerebellar ataxia type 7 (SCA7) is caused by a mutation in the *ATXN7* gene. We focused on RNAs with at least 10 CAG repeats and identified several hundred mRNAs, long non-coding RNAs (lncRNAs), and circular RNAs (circRNAs) originating from 49 genomic *loci*. These *loci* differ in the rate of length polymorphism in the population, and some of the tracts show an interesting pattern of adjacent CAA triplet, as well as present variable interruption profiles. For *ATXN7 locus*, we investigated two circRNAs for which we confirmed the expression in the human brain, SCA7 fibroblasts and blood samples. Moreover, we report an intriguing example of circRNA containing a mutant CAG tract, experimentally validated in SCA7 patients.

## Introduction

CAG repeat tracts are examples of short tandem repeats (STRs) present in the human genome. CAG repeats are among triplet repeat tracts overrepresented in the transcriptome and are present in protein-coding and non-coding RNAs (ncRNAs) [1,2]. The length of CAG tracts generally increases with the complexity of organisms and has been associated with better brain functioning [3,4]. On the other hand, expansions of these sequences above the defined threshold have adverse effects and cause neurodegenerative disorders, including polyglutamine (polyQ) diseases, like Huntington’s disease (HD) and several types of spinocerebellar ataxias (SCAs). While the significance of CAG repeats in protein-coding sequences is well established [5,6], their prevalence and potential functional roles in non-coding RNAs remain underexplored. Various ncRNAs, including long non-coding RNAs (lncRNAs) and circular RNAs (circRNAs), emerge as key players in diverse cellular processes, often in a cell-type- and developmental stage-dependent manner, which causes additional challenges in studying them [7–9]. lncRNAs are a highly heterogeneous group involved in chromatin remodeling, transcriptional regulation, and RNA processing. circRNAs, in turn, are covalently closed transcripts with high stability and often exhibit tissue-specific expression and regulatory potential, particularly in the nervous system [7,8].

## Materials and methods

Human RNAs containing at least 10 uninterrupted CAG repeats were retrieved from GENCODE release 39 [10] for mRNAs and lncRNAs, and from circAtlas 3.0 [11] for circRNA, as described before [2]. Human fibroblasts, SCA7 (GM03561) and healthy controls (GM07525, GM04503) (Coriell Institute) were cultured in recommended conditions. Human blood samples were collected from the three symptomatic patients with SCA7 belonging to Polish Family 1 [12] and healthy individuals, using PAXgene Blood RNA Tubes (Qiagen). Total RNA from the human brain and liver was purchased (ThermoFisher). Total RNA was isolated from fibroblasts with TRI reagent (Invitrogen) using Total RNA Zol-Out D kit (A&A Biotechnology) and from blood using PAXgene Blood RNA Kit (Qiagen). RNA concentration and quality were determined using a DeNovix spectrophotometer or Qubit fluorometer and TapeStation (Agilent). cDNA was generated using SuperScript IV Reverse Transcriptase (Invitrogen) with random primers. RT-qPCR assays were run using SsoAdvanced Universal SYBR Green Supermix (BioRad) on a CFX Connect (BioRad). RT-PCRs were run using GoTaq Flexi DNA Polymerase (Promega) on a T100 Thermal Cycler (BioRad), the products were loaded on an agarose gel, purified and sequenced. Primer sequences are listed in Table S1. More details on methods are provided in the Supplementary Text. Statistical analyses were performed using GraphPad Prism.

## Results

In this study, we focused on RNAs with at least 10 CAG repeats, as such sequences are more likely to have a variable length in the population and/or to fulfill a functional role in the molecule. We identified human RNAs containing at least 10 CAG repeats in the reference datasets: 135 mRNAs, 27 lncRNAs, and 93 circRNAs (Fig. S1, Table S2) originating from 49 genomic *loci*. Specifically, the identified mRNA, lncRNA, and circRNA sequences originated from 35, 11, and 20 genomic *loci*, respectively (Fig. 1A), with a very variable number of different transcript variants for each *locus* (Fig. S1). Identified *loci* include genes involved in transcription (*E2F4, IRF2BPL*), Alu-mediated gene expression regulation (*MIR205HG*), neuronal function (*HTT, ATXN2*), and chromatin remodeling (*SMARCA2, EP400*). Several identified transcripts correspond to genes for which the CAG tract mutations have been described, mainly genes implicated in polyQ diseases. From most identified *loci* (exactly 35), mRNAs are produced, indicating that ≥ 10 CAG repeat sequences are predominantly associated with protein-coding genes. A substantial number of *loci* (16) produce both mRNAs and circRNAs, suggesting that many CAG-containing exons in mRNAs undergo back-splicing, producing circular isoforms. In contrast, fewer CAG repeat-bearing transcripts (11) are classified as lncRNAs-producing *loci*, with only one overlapping with *loci* producing circRNAs (Fig.1A).

**Figure 1.**
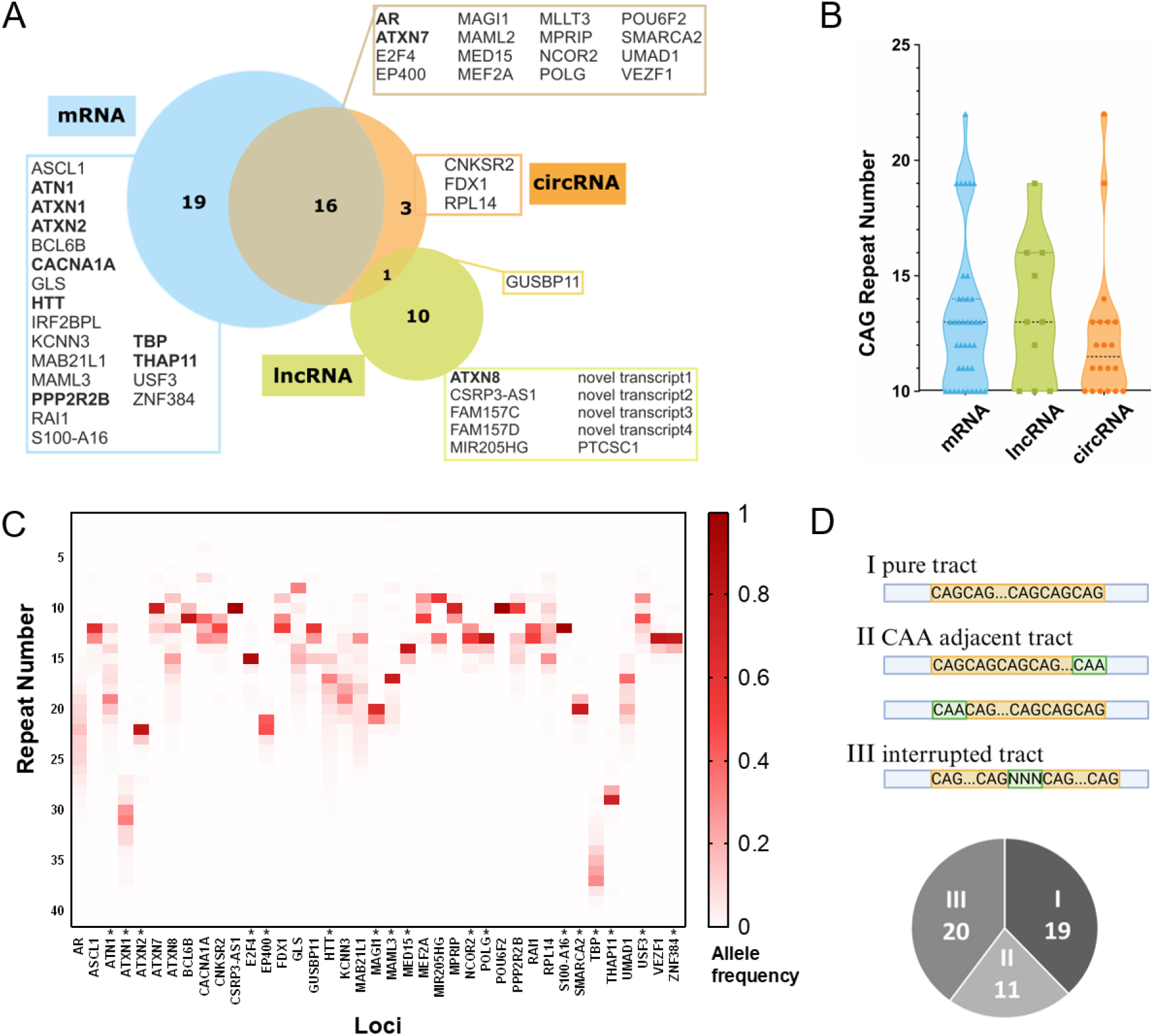
RNAs with at least 10 CAG repeats: biotypes, tract polymorphism, and sequence patterns. **A** Venn diagram of 49 genomic *loci* with ≥10 CAG repeats, showing the classification of molecules: mRNA (blue), lncRNA (green), and circRNA (orange), with indicated *loci* that produce two different types of RNAs. Bolded gene names represent *loci* known to be associated with repeat expansion disorders. novel transcript1 - ENST00000669801.1; novel transcript2 - ENST00000649462.1; novel transcript3 - ENST00000469183.5; novel transcript4 - ENST00000244820.2/LOC93463. **B** Distribution of CAG tract lengths in different RNA biotypes (based on reference sequence in databases). Violin plots show kernel density estimation of the distribution of CAG tract length in mRNAs (blue), lncRNAs (green), and circRNAs (orange). Each dot represents an individual RNA molecule. Horizontal dashed lines within the violins represent the median values. **C** Heatmap showing allele frequency distributions of repeat lengths across CAG-containing *loci*, based on data deposited in TR-Atlas (338,963 human samples in diverse ancestries; data were available for 39 of the analyzed *loci*). Repeat number refers also to other triplets, as not all tracts are pure CAG (loci with interruption are marked with an asterisk). **D** Classification of CAG tract types and pie chart with the number of *loci* in each category. All tracts contained at least 10 CAG repeats and were classified as: (I) pure, (II) with adjacent CAA triplet (at 3’ or 5’ site), or (III) interrupted (containing in the tract of CAG units at least one trinucleotide different from CAG). A total of 50 tracts were classified due to the presence of a tract that has both a CAA adjacent codon and an interruption in the *MAML2* loci.

Regarding CAG tract lengths in the reference sequences in databases, most identified RNAs contained 10–15 CAG repeats, though longer tracts were observed in all three RNA classes (Fig. 1B). Recent advancements in the analysis of polymorphism of repeat tracts in population [13,14] enable us to analyze selected CAG-containing *loci* in this aspect. Due to sequence/structure-related aspects and/or functional pressure on specific repeat numbers, the CAG tracts at individual *loci* vary significantly in terms of the degree of length variation in the population. Some of the *loci*, like *CSRP3-AS1, POU6F2*, and S100-A16, show the dominance of a single-length variant, whereas other loci, like AR, HTT, and MAB21L1, are characterized by a large variation of the number of triplet units in the analyzed population [14] (Fig. 1C).

We classified the CAG repeat tracts in identified *loci* as: (I) pure, (II) with adjacent CAA, or (III) interrupted (Fig. 1D). Many *loci* (19), such as *ATXN7, CACNA1A*, and *PPP2R2B*, contain uninterrupted CAG repeats. These pure tracts are typically more prone to expansion, increasing the risk of disorders like spinocerebellar ataxias (SCAs). Other *loci* (20) contain interruptions within the CAG repeat, which can be single, like for *ATXN2* and *HTT*, or multiple, like for *SMARCA2* and *TBP*. These interruptions can influence repeat stability, often reducing expansion rates and potentially delaying disease onset [15,16]. Some *loci* (11), including *AR* and *MIR205HG*, have CAA triplets adjacent to CAG tracts that may also contribute to repeat stability. CAA, like CAG, encodes glutamine, therefore leading to a longer polyQ tract in case of translation.

For the selected *ATXN7 locus*, we identified nine circRNAs that contained a CAG repeat tract (Figure S2). The expansion of the CAG tract in the *ATXN7* gene encoding the ataxin-7 protein (which contains an abnormal polyQ tract when mutated) causes SCA type 7 (SCA7) [6]. CircRNAs identified at the *ATXN7* locus were formed from various exon combinations, ranged in length from 336 to 1106 nts, and all contained complete exon 3 of *ATXN7* mRNA with a binding site for NF-kappaB p65 subunit [17] (Fig. S2). For further experimental validation, we selected hsa-ATXN7_0001 (circATXN7(3,4).1; named later on “circ1”) and hsa-ATXN7_0005 (circATXN7(2,3,4,5).1; named “circ5”), based on the substantial expression levels of these circRNAs in human brain and blood [11]. We verified the sequence of the BSJ site of these circRNAs in human fibroblasts and brain tissue (Fig. 2A). Next, we analyzed circ1, circ5, and linear *ATXN7* RNA levels in human fibroblasts. We observed reduced expression of circ1 and circ5 in SCA7 fibroblasts, to 60-70% of the level in the control cell lines (Fig. 2B, upper graph). The linear *ATXN7* RNA had a similar level in the SCA7 line compared to the control lines. As a consequence, a lower circular to linear ratio was observed in the SCA7 line for both circRNAs (a tendency for circ1 and statistically significant result for circ5) (Fig. 2B, lower graph), suggesting that biogenesis of these circRNAs may be altered in SCA7. For circ1, we were able to design an RT-PCR that spanned both the CAG tract and the BSJ site. Our analysis, confirmed by Sanger sequencing, revealed the presence of an expanded CAG tract (with at least 56 repeats) within circ1 in RNA isolated from SCA7 fibroblasts (Fig. 2C). We also analyzed blood samples of three symptomatic SCA7 patients [12] and found no apparent differences in the levels of circ1, circ5, or linear *ATXN7* RNA, as compared to healthy individuals’ samples (Fig. 2D). We validated that, also in the blood of SCA7 patients, circ1 with mutant CAG tract (with at least 42 repeats) is present (Fig. 2E). Additional features of circ1 expression predispose it for further studying: (I) in blood samples circ1 product amplified much earlier than circ5, (II) circ1 showed higher expression of circular form in the brain as compared to the liver, while the linear form of *ATXN7* RNA showed a slightly lower level in the brain than in the liver (Fig. 2F). We also examined the effect of repeat tract expansion on the modelled secondary structure of circ1 and found no apparent differences between circ1 with 10 or 70 CAG repeats in the regions spanning the BSJ site, as well as the p65 binding motif was still present in both structures (Fig. 2G). As expected, the expanded CAG tract formed an elongated hairpin structure, the presence of which could be expected to cause some adverse effects due to RNA toxicity pathways [18].

**Figure 2.**
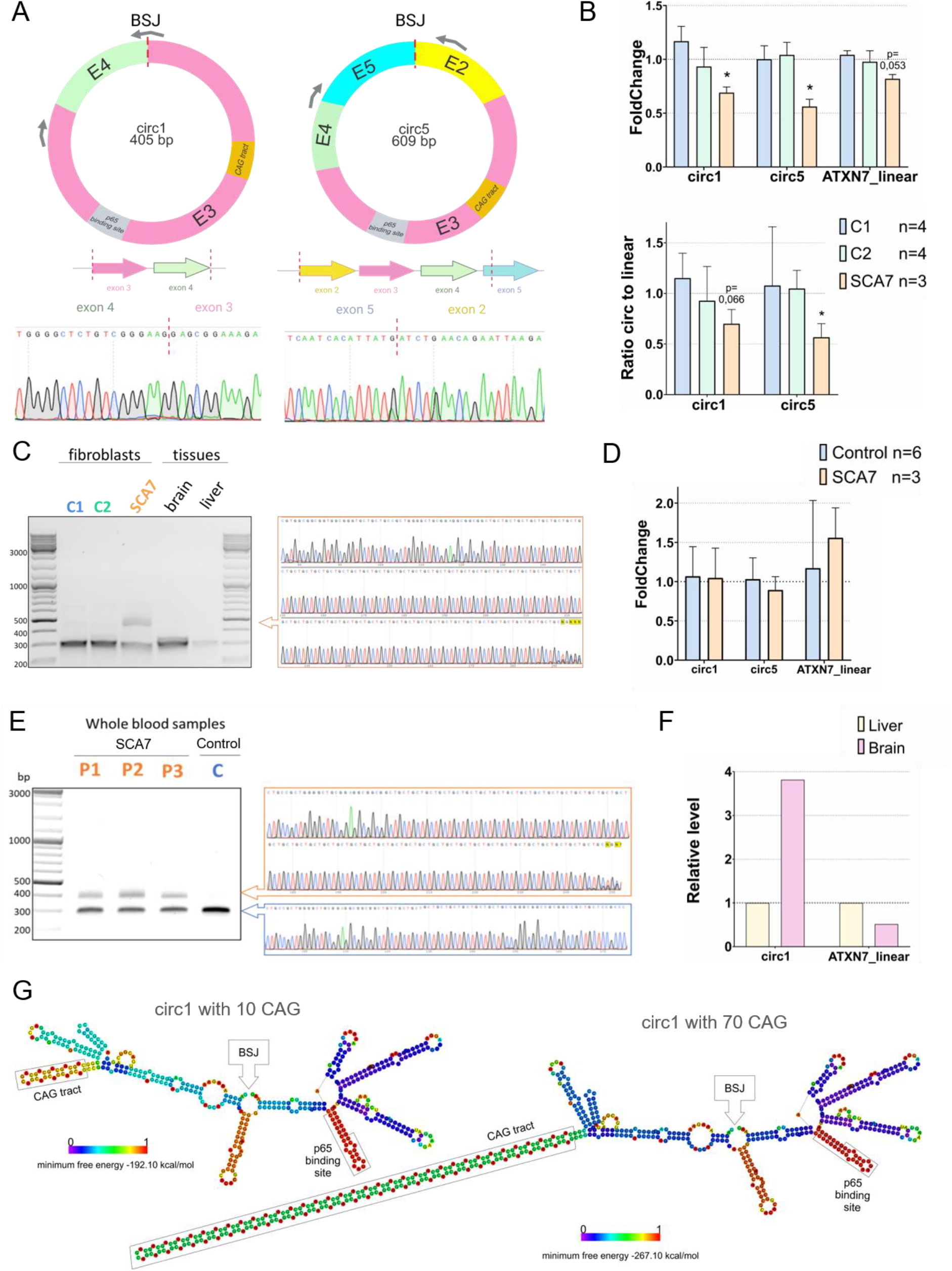
Experimental validation of *ATXN7* circRNAs and mutant variant identification in SCA7 cells and patients’ blood. **A** Schematic representation of circ1 (hsa-ATXN7_0001/Uniform ID: circATXN7(3,4).1) (left panel) and circ5 (hsa-ATXN7_0005/Uniform ID: circATXN7(2,3,4,5).1) (right panel) with exon composition below and Sanger sequencing results showing back-splice junction (BSJ), indicated by dashed red line. Grey arrows indicate the location of primers used for BSJ site sequencing. **B** The relative level of circ1, circ5, and linear *ATXN7* mRNA in human fibroblasts was assessed using RT-qPCR. C1 (n=4), C2 (n=4) - healthy individuals; SCA7 (n=3) - patient’s line, where n is the number of independent sample collections. For circ1 and circ5 expression analysis, we used primers that spanned the specific BSJ sites (but not the CAG tract) and obtained single products of the expected length. The mean result from the control lines was set as 1 for normalization. As a reference, expression levels of *HIPK3* (circHIPK3(2).1) and *N4BP2L2* (circN4BP2L2(3,4,5S,S6).2) were used for circRNA, and *GAPDH* and *EEF2* for linear RNA. The circ/linear RNA ratio was calculated using linear RNA Ct as a reference; values are presented as fold changes over linear RNA. Statistical significance was determined by a one-sample t-test with hypothetical mean = 1. **C** RT-PCR products (containing CAG repeat tract and BSJ site) for *circ1* obtained from RNA isolated from human fibroblasts (C1, C2 - healthy individuals; SCA7 - patient’s line) and human tissue lysates. The slower migrating band obtained for the SCA7 line was sequenced and confirmed to contain an expanded CAG tract. **D** The relative level of circ1 and circ5 was assessed using RT-qPCR in human whole blood samples from healthy individuals (control, n=6) and SCA7 patients (n=3), where n is the number of individuals. For normalization, the mean result from control lines was set as 1. As a reference, expression levels of *HIPK3* and *N4BP2L2* were used for circRNA, and *GAPDH* and *EEF2* for linear RNA. Statistical significance was determined by an unpaired t-test. **E** RT-PCR products of circ1 containing CAG repeat tract and BSJ site, obtained from human blood samples (C - healthy individual; P1, P2, P3 - SCA7 patients’ samples). Indicated products were sequenced. The slower migrating bands were confirmed to contain an expanded CAG tract. **F** The relative level of circ1 in human tissue lysates was assessed using RT-qPCR (n=1). For normalization, the mean results from liver tissue were set as 1 and used as a reference for brain tissue; *GAPDH* and *EEF2* were used as a reference for linear RNA. **G** Predicted secondary structures formed by *circ1* with 10 CAG (wild-type variant) and 70 CAG (mutation in SCA7) using RNAfold. The CAG region, BSJ site, and p65 binding motif are indicated in both structures.

## Discussion

As ncRNAs are generally known to have higher expression and specified functionality in the brain, their investigation is an emerging field in molecular neurobiology. Also, for neurodegenerative diseases with a crucial protein component identified, it becomes essential to investigate ncRNAs that may significantly contribute to altered pathways and serve as biomarkers, due to their presence in EVs and/or high stability, like in the case of circRNAs [19]. One example is a recently described circRNA from the *HTT locus*, but not containing CAG repeats, that was described as a modifier of HD phenotype [20].

The investigated circRNAs from the *ATXN7 locus* are predicted to bind miRNAs and RBPs [11], which needs further validation. Interestingly, circRNA-forming exon 3 of *ATXN7* that contains a CAG tract also has an experimentally validated binding site for the p65 subunit of NF-kappaB (Fig. 2G). In case of increased levels of this circRNA in cancer, a sequestration of p65 in cytoplasm and a lack of proper functioning of NF-kappaB in transcriptional regulation in the nucleus were shown [17]. Moreover, NF-kappaB dysfunction was already described in SCA7 but explained as caused by inhibition of proteasome activity by mutant ataxin-7 [21]. Whether additional pathways may disrupt NF-kappaB functioning in SCA7 needs further investigation.

To the best of our knowledge, the presence of the CAG tract has not been explored yet in circRNAs, including the *ATXN7 locus*. Generally, the CAG tract length in this *locus* is not highly polymorphic in the population [13,14] as ~75% people have 10 CAG repeats in *ATXN7*, the second frequent variant is 12 CAGs (~12%) [14] (Fig. 1C). Other identified circRNAs have longer and or more variable CAG tracts (Table S2), which may suggest the functionality of these tracts in circRNAs. From our initial experiments, especially circ1 is a good candidate for further investigation, due to its higher expression levels in human blood and brain tissue (as compared to circ5), and also, for this circRNA, we were able to confirm the presence of a mutant variant in both SCA7 patients’ fibroblasts and blood. This points to further investigation of this circRNA, e.g., in extracellular vesicles (EVs) isolated from blood or specific brain cell types.

## Acknowledgments

We are grateful to patients and healthy individuals for consenting to use their samples. We acknowledge the use of the infrastructure developed under the project NEBI - National Research Center for Imaging in the Biological and Biomedical Sciences, POIR.04.02.00-00-C004/19, co-financed through the European Regional Development Fund (ERDF) in the frame of Smart Growth Operational Programme 2014-2020 (Measure 4.2 Development of modern research infrastructure of the science sector). We acknowledge the Laboratory of Single Cell Analysis and the Laboratory of Genomics ICHB PAS for access to equipment. We would like to thank the lab members for the discussions, as well as Paweł and Piotr Świtonski for their support of this study. Figures were created using biorender.com and GraphPad Prism.

## Author Contribution Statement

Conceptualization: AF, WP, MJC, MWW. Experiments: WP, MWW. Data analysis: WP, AF, MWW, MJC, EK, TW. Human samples collection: JD, JS. Funding acquisition and supervision: AF. Manuscript draft and figures: WP, AF. All authors reviewed and approved the final manuscript.

## Ethical Approval

The research experiment received the approval of the Bioethics Committee at the Medical University in Gdańsk (NKBBN/462/2020 and KB/462-48/2025). Informed consent was obtained from all individual participants included in the study.

---

**Supplementary Figure 1.**
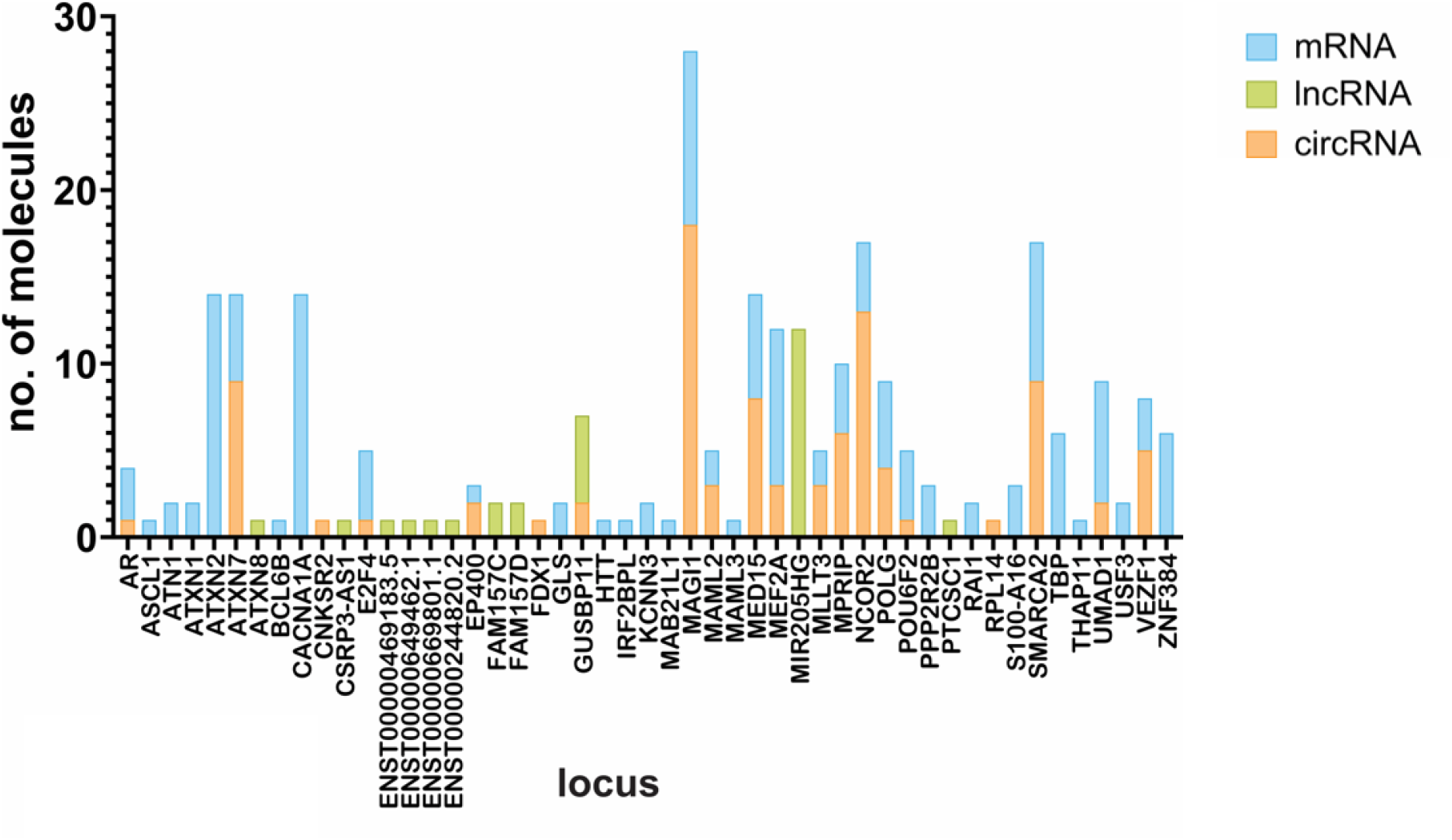
The number of variants of CAG-containing RNA molecules across genomic *loci* and RNA biotypes. The number of mRNA (blue), lncRNA (green), and circRNA (orange) molecules with CAG tracts detected at each genomic *locus* is presented.

**Supplementary Figure 2.**
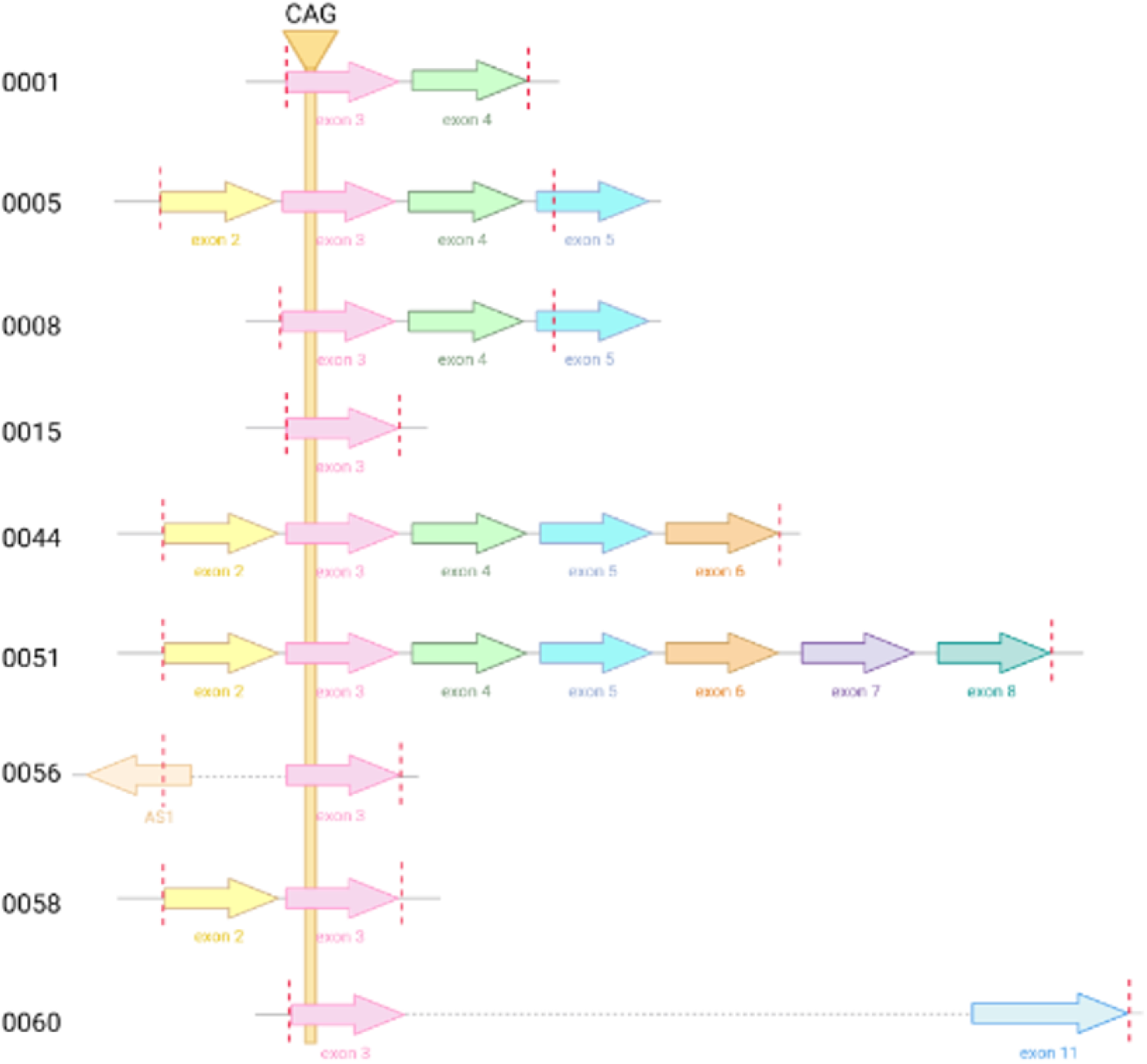
Graphic representation of exon and intron composition of CAG repeat-containing circRNA molecules from the *ATXN7 locus*. The dashed red line indicates the back-splice junction site.

**Supplementary Figure 3.**
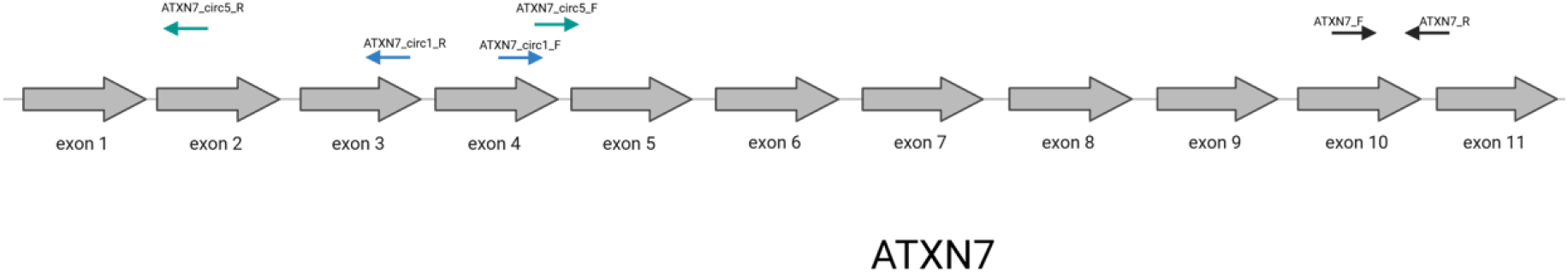
Graphical representation of exons in the *ATXN7 locus*. Arrows indicate the locations of primers used for expression quantification: blue for circ1, green for circ5, and black for the linear transcript.

**Supplementary Table 1.**
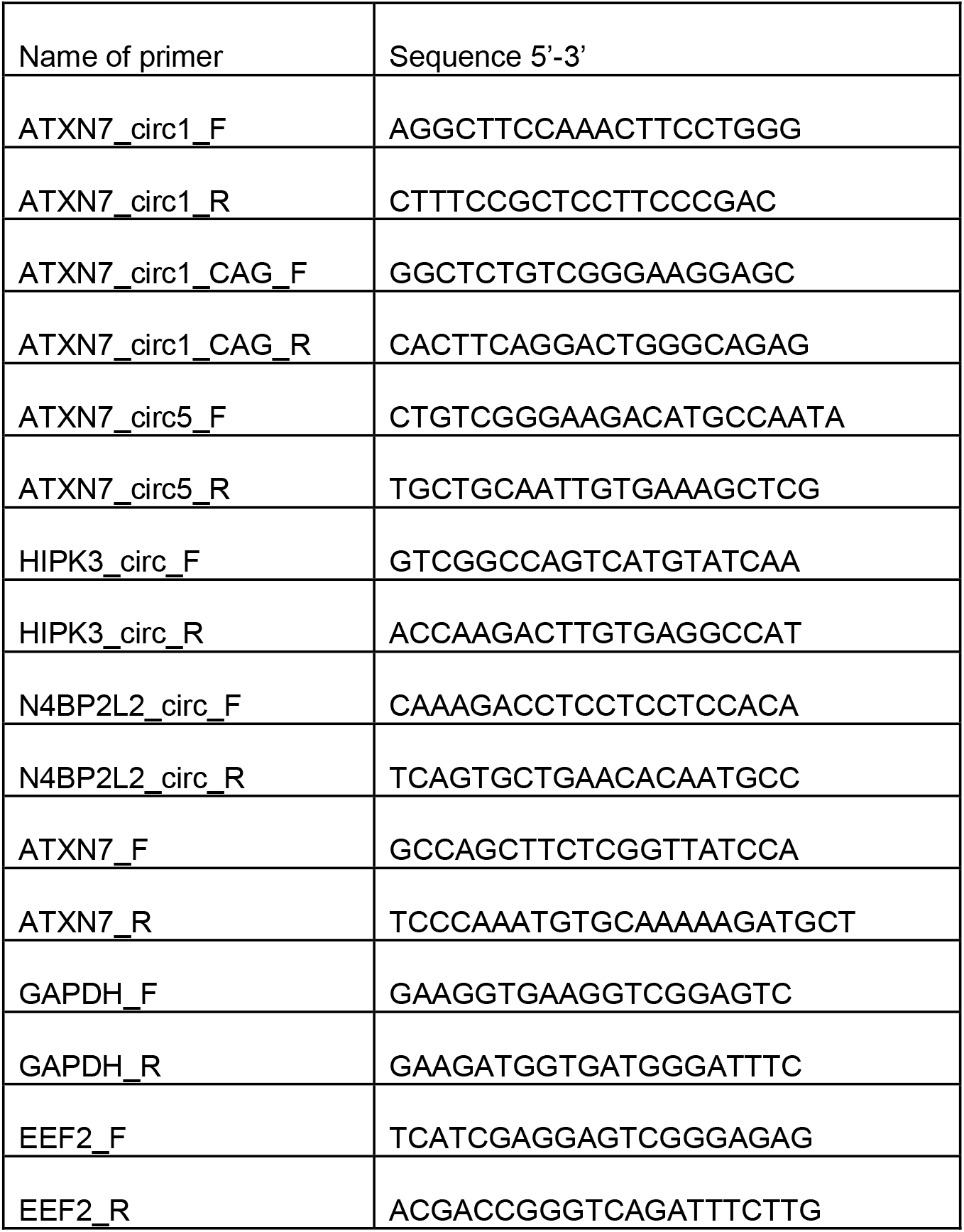
Sequences of primers used in PCR.

**Supplementary Table 2**. Lists with information about identified mRNAs, lncRNAs, and circRNAs containing a CAG tract composed of at least 10 units.

## Supplementary Text

### Detailed Materials and Methods

#### Retrieval of coding and ncRNAs with CAG tracts from databases

An in-house Python script was used to identify human RNAs containing at least 10 CAG repeats among protein-coding transcripts, lncRNAs, and circRNAs, as previously described [2]. Protein-coding transcripts and lncRNAs were retrieved from GENCODE release 39, and human circular RNA datasets from circAtlas 3.0. Identified RNAs containing CAG tracts are listed in Table S2.

#### Cell lines and human samples

Fibroblast cell lines — human SCA7 (GM03561) and two healthy controls (GM07525, GM04503) (Coriell Institute) — were cultured in MEM (Gibco) supplemented with 10% fetal bovine serum (EurX), 100 U/ml penicillin–streptomycin (Gibco), and 2 mM L-glutamine (Gibco), and passaged using trypsin. Total RNA was isolated from cell pellets with TRI Reagent (Invitrogen) using the Total RNA Zol-Out D kit (A&A Biotechnology). Briefly, cell pellets suspended in TRI Reagent were mixed with one volume of 100% ethanol and loaded onto microcolumns. Following RNA binding, on-column DNase digestion was performed for 30 min at 37 °C. The isolation was performed following the manufacturer’s protocol. RNA was eluted in RNase-free water after a short incubation at room temperature. RNA concentration was determined using a spectrophotometer (DeNovix).

Total RNA from human brain (First Choice Human Brain Total RNA) and liver (First Choice Human Liver Total RNA) was purchased from ThermoFisher.

Human blood samples were collected from three symptomatic patients with SCA7 belonging to Polish Family 1 [12] and from healthy individuals, using PAXgene Blood RNA Tubes (Qiagen). Total RNA was isolated from peripheral whole blood using the manual protocol of the PAXgene Blood RNA Kit (PreAnalytiX), according to the manufacturer’s instructions. Following collection, tubes were incubated for 2 h at room temperature to ensure complete lysis of blood cells and stored at –80 °C until extraction. Briefly, frozen samples were thawed and centrifuged, the supernatant was removed, and the pellet was washed with RNase-free water. After a second centrifugation, the pellet was resuspended and incubated in buffer with proteinase K. Lysates were homogenized by centrifugation through the PAXgene Shredder spin column, and the cleared flow-through was mixed with ethanol and loaded onto PAXgene RNA spin columns. After binding, RNA was purified through sequential washes. Between the first and second wash steps, the membrane was treated with DNase I to remove trace DNA. RNA was eluted in elution buffer, heat-denatured, and quantified using a Qubit 4 fluorometer with the Qubit RNA BR Assay kit (Invitrogen). RNA integrity was assessed using the 4150 TapeStation system (Agilent).

### RNA isolation, reverse transcription, and RT-qPCR

cDNA was synthesized from 0.5 µg RNA using SuperScript IV Reverse Transcriptase (Invitrogen) with random primers, according to the manufacturer’s protocol. The resulting cDNA was diluted 1:10, and 2.5 µl was used per RT-qPCR reaction. Reactions were prepared using the SsoAdvanced Universal SYBR Green Supermix (BioRad) and run on a CFX Connect system (BioRad). Each RNA sample was assayed in triplicate. Statistical analyses were performed in GraphPad Prism.

#### Primer design

Primers were designed using the NCBI Primer-BLAST tool (https://www.ncbi.nlm.nih.gov/tools/primer-blast). Standard parameters (58 °C < Tm < 62 °C, amplicon sizes between 90–250 bp) were applied. Binding sites for primers at the *ATXN7 locus* are shown in Figure 2A and Figure S3. All primers were tested for efficiency and specificity of amplification before use. Primer sequences are listed in Table S1.

#### Back-splice junction confirmation for circRNAs

RT-PCRs were performed using GoTaq Flexi DNA Polymerase (Promega) on a T100 Thermal Cycler (BioRad). Products were resolved on 1.5% agarose gels, purified using PureLink Quick Gel Extraction Kit (Invitrogen), and sequenced.

